# Somatic maintenance impacts the evolution of mutation rate

**DOI:** 10.1101/181065

**Authors:** Andrii Rozhok, James DeGregori

## Abstract

The evolution of multi-cellular animals has produced a conspicuous trend toward increased body size. This trend has introduced at least two novel problems: an elevated risk of somatic disorders, such as cancer, and declining evolvability due to reduced population size, lower reproduction rate and extended generation time. Low population size is widely recognized to explain the high mutation rates in animals by limiting the presumed universally negative selection acting on mutation rates. Here, we present evidence from stochastic modeling that the direction and strength of selection acting on mutation rates is highly dependent on the evolution of somatic maintenance, and thus longevity, which modulates the cost of somatic mutations. We argue that this mechanism may have been critical in facilitating animal evolution.

## Introduction

Increasing body size has been one of the major trends in animal evolution across many taxa, as formulated in Cope’s rule [1,2]. The evolution of larger bodies, particularly for land animals and especially for mammals, introduces some fundamentally new evolutionary challenges. The carrying capacity of ecosystems limits biomass per group/species, so larger body size leads to reduced population size. Furthermore, large animals generally demonstrate lower numbers of progeny, longer generation times, and more advanced ages of first reproduction [3,4]. In aggregate, such changes weaken selection that can act on a population and thus should negatively affect evolvability. This general reduction in evolvability should, however, be at least partially alleviated by diversity facilitated by sexual reproduction.

The mutation rate (MR) is another critical evolvability parameter [5]. It is believed that selection generally acts to lower MR [6–8], and the significantly higher MRs observed in animals compared to unicellular organisms have been argued to result from the reduced power of selection imposed by small population sizes [9–11]. Germline (gMR) and somatic (sMR) mutation rates are linked, as they employ the same basic DNA replication and repair machinery [12–14]. While elevated gMR improves evolvability, the ensuing higher sMR should elevate the risk of somatic disorders, such as cancer [15]. For cancer, increasing body size is expected to increase the frequency of oncogenic mutations by increasing the number of target cells [16]. Somatic mutations also contribute to aging and a variety of aging-related diseases [17]. The increased cost of sMR should thus exert negative selective pressure on gMR in larger animals.

Recent evidence demonstrates that the sMR in some animal tissues can be significantly higher than the rate inferred from observed mutations, because somatic purifying selection is very effective at eliminating damaged somatic cells [18]. Many mechanisms, such as various tumor suppressor gene functions (including DNA damage induced apoptosis) [19], autophagy [20], purifying somatic selection [18,21], and immune surveillance [22], should buffer the costs of somatic mutations and in aggregate promote lifespan extension by maintaining tissue integrity. We will collectively call these mechanisms - the *somatic maintenance program* (SMP).

We present theoretical evidence from Monte Carlo modeling indicating that somatic maintenance not only improves individuals’ survival for large animals by reducing sMR costs, but should have played a crucial role in animal evolution by impacting the evolution of gMR. We show that positive selection for increased body size promotes positive selection for extended longevity by improving SMP. Our results also indicate that positive selection acting on traits that do not impact somatic risks also promotes selection for an improved SMP. In both cases, evolution toward increased gMR was observed because of the reduced sMR cost, which dramatically improved evolvability of the simulated population.

## Results

### Theoretical introduction to the modeling

We built a stochastic model of evolution in animal populations, incorporating reproduction and survival (**Fig. 1**), whereby each individual’s trait is inherited with variance proportional to gMR (for code, see **Supplements: Section 1a**). Traits are assumed to be polygenic and exhibit phenotypic variation in the population. In particular, MR is assumed to be a highly polygenic trait, given the many genes responsible for DNA repair, DNA replication, damage avoidance (e.g. anti-oxidant defenses), and mutagen detoxification, which in aggregate can determine MR. The evolution of body size, somatic maintenance and germline mutation rate was then tracked under various regimens of selection (see also **Methods: Model algorithm**).

**Fig. 1.**
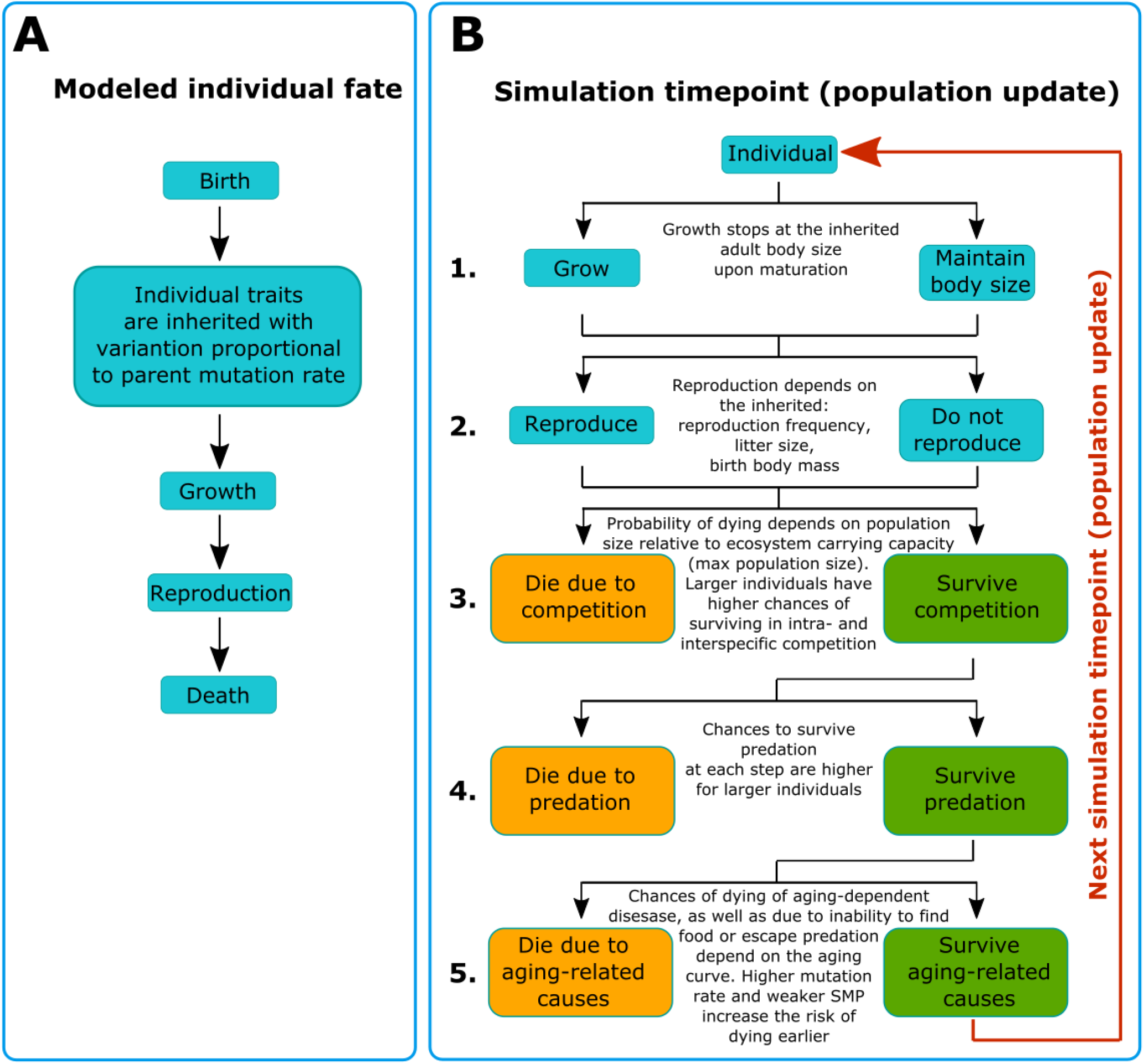
A scheme of the model simulations. (**A**) Stages of an individual simulated lifespan. (**B**) At each timepoint during the simulation the modeled population undergoes 5 main updates: 1. Individuals that have not reached maturity increase their body mass following their growth curve, starting from the initial birth mass and up until they reach their inherited body mass (parent body mass with variation proportional to parent mutation rate); mature individuals remain at the same body mass. 2. Each individual past maturation reproduces with a certain inherited frequency of reproduction, producing on average an inherited number of progeny per litter, each progeny’s birth body mass is inherited from its parent with variation proportional to the parent’s mutation rate. Each individual is tried in a binomial trial with a small probability (at each timepoint) of dying of three main causes: 3. Death following limitations imposed by ecosystem carrying capacity which allows for a certain maximum population size and promotes intra-specific and inter-specific competition for resources if population numbers exceed this capacity. 4. Death caused by predation is modelled based on the Lotka-Volterra model of predator-prey interaction (*34, 35*). 5. Death caused by physiological aging, such as due to cancer, frailty or other age-related causes; the probability is negligible early in life but increases exponentially with age; the speed of increase of the probability of death caused by aging depends on an individual’s aging profile which is determined by the aging curve as explained in **Fig. 3A**, “Theoretical introduction to the modeling” subsection of **Results** and “The somatic maintenance program paradigm” subsection of **Methods**.

The model should reasonably approximate a sexually reproducing population. The model operates with single-parent reproduction model so that each individual descends from one parent. In this regard, technically it is tempting to view it as a model of an asexual population. However, the fundamental differences that lie between sexual and asexual populations (aside from the issue of purging deleterious mutations) are the amount of variation produced per the same sized population per generation and the presence of genetic recombination. Variance of inheritance in our model is too high to be assumed as having been generated by mutations accumulating along a clonal lineage and equals 10% of a trait’s value per generation within 95 percentile. Genetic recombination associated with sexual reproduction segregates different genes and thus obstructs genetic linkage characteristic of inheritance during clonal reproduction. However, as **Fig. 2** demonstrates, the role of genetic recombination in the evolution of multi-genic traits is inversely proportional to the number of genes and alleles encoding a trait. The gene pool shown in **Fig. 2A** encodes a multi-genic trait, whose distribution in a population is shown in **Fig. 2C**. As can be seen in **Fig. 2C**, an increasing number of genes encoding a trait decreases the variance of the encoded trait when randomly sampled from a population. This means that randomizing factors, such as genetic recombination, that randomly shuffle alleles for each gene in a population will have a smaller effect on the phenotypic expression of a trait encoded by many genes compared to oligo- and monogenic traits. Multi-genic traits also provide multiple combinations of alleles for each of the encoding genes that can result in the same net expression of the phenotype. Based on this logic and simulation, mutator phenotypes, given the highly polygenic nature of mutation rate, will not be easily segregated from adaptive traits by genetic recombination as occurs with monogenic traits. We therefore did not simulate the processes of allelic segregation by recombination in order to reconstruct a sexual population (which would require many unwarranted assumptions), but modeled the distribution of mutation rate that was independently inherited from other traits and was not directly subjected to selection in the model (**Fig. 2D**). As such, the model only operates with the net ultimate change of a trait over generations. The assumed multigenic nature of the simulated traits also means that both segregation of alleles by recombination and aggregation of alleles by co-selection are impeded. We therefore assume that the net co-evolution of a pair of multigenic traits will ultimately depend on selection acting on these traits that can overcome the effects of genetic recombination.

**Fig. 2.**
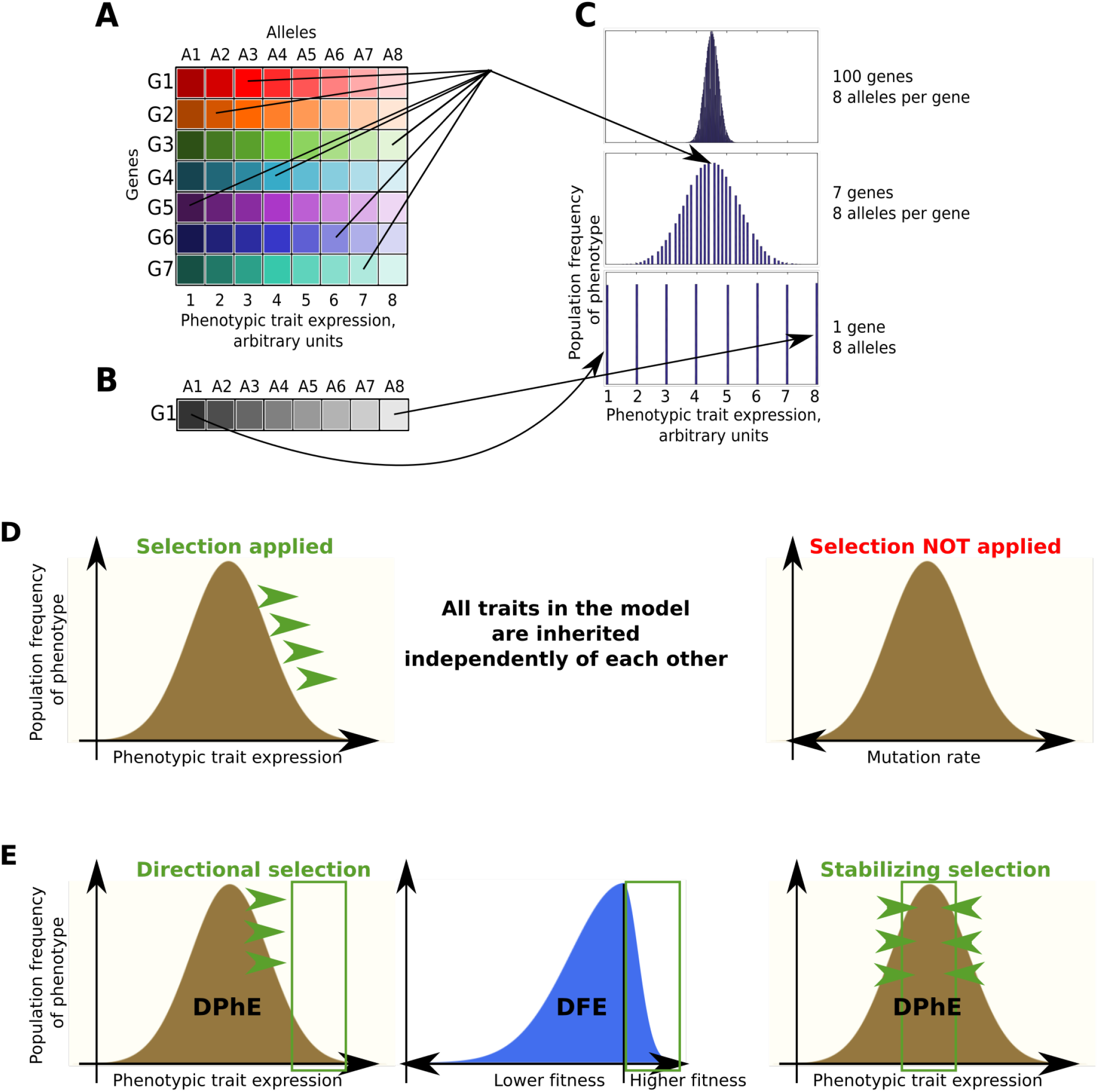
Multigenic inheritance alters the role of linkage disequilibrium in the evolution of phenotypic traits. (A) The color table illustrates a simplified model of a population’s gene pool for a trait encoded by multiple genes. For simplicity, all the 7 genes are represented in a population by 8 alleles for each gene. Each allele differentially impacts the overall expression of a trait (shown in arbitrary units on the lower X-axis). (B) A population’s gene pool for a trait encoded by one gene with 8 alleles which impact the phenotypic trait expression as shown in (A). (C) A Matlab-generated (see code in **Supplemental Materials Section 6**) distribution of phenotype expression in a population of 100,000 individuals with different number of genes encoding the trait and each represented by 8 alleles. The X-axis in all the three charts is the same from 1 to 8, the Y-axis is population frequency of the phenotype (no scale for simplicity). Arrows demonstrate the allelic composition of phenotype examples generated based on multi-genic (panel A) and monogenic (panel B) trait inheritance. The probability that all alleles in a multigenic trait will change the trait in the same direction (for instance all A8 alleles in panel A) is inversely proportional to the number of genes and is the product of the probabilities of each particular allele occurring in the given phenotype. An increased number of genes affecting the trait, therefore, decreases the likelihood that a random sampling event, such as genetic recombination, will result in a significant deviation of the resulting trait expression. (D) Assuming the multigenic nature of mutations rates and body size (primary modeled traits), we modeled both as normally distributed (see panel C) and independently inherited. Selection was applied on body size (or other modeled trait) but <u>not</u> on mutation rate. The evolution of mutation rate in the model was completely independent of other traits. (E) The left and right charts demonstrate a typical distribution of phenotypic trait expression (DPhE) in a population. Directional selection (left chart, green arrows) will favor extreme phenotypes from the target phenotype tail (left chart, green box). Stabilizing selection (right chart, green arrows) will favor the population mean which is the target phenotype (right chart, green box). Under both scenarios, most germline mutations will be in the negative (disadvantageous) tail of the distribution of fitness effects, DFE (middle chart), imposing fitness a cost for germline mutations. The exact ratio of the tails of the resulting DFEs (and thus the cost/benefit ratio of germline mutations) are determined by the mode and strength of selection acting on a phenotype.

While we explicitly model the cost of sMR realized in reduced body fitness, our model does not incorporate lethal germline mutations, as they affect the population randomly (all zygotes have roughly the same chance of acquiring a lethal mutation) and thus are irrelevant to selection. Our model operates with *distributions of phenotypic effects, DPhE,* of germline mutations, which in general can be safely modeled as normally distributed in accordance with the Shelford law of tolerance [23], and the cost of germline mutations (which relates to the *distribution of fitness effects, DFE*) depends on and is imposed by the mode of selection applied to a trait as shown in Fig. 2E. For instance, under stabilizing selection (population mean trait is favored) a high germline mutation rate which produces more widely distributed trait values (wider DPhE) will impose a fitness cost on mutations with large phenotypic effects (deviating from the mean). Under positive selection, respectively, the positively selected tail of the DPhE (deviant trait values) will be favored. However, under both regimens the vast majority of phenotypically expressed germline mutations will lower fitness and be in the negative tail of the DFE (Fig. 2E). Our model for the cost of germline mutations therefore avoids artificial assumptions and replicates the natural mechanism for how selection determines the fitness cost of germline mutations. Noteworthy, this cost will differ for different traits depending on the current DPhE of the trait in the population at any given moment of evolution, as well as the current mode and strength of selection acting on the trait.

The model incorporates three major factors of mortality, including aging. Human life tables indicate that aging proceeds exponentially, whereby mortality and diseases accelerate at advanced ages (e.g. https://www.ssa.gov, https://seer.cancer.gov). The combined action of SMP mechanisms provides for an extended early period of high body fitness with little to no decline. We generalized this complex program in a curve that describes modeled animal mortality of physiological causes schematically shown in **Fig. 3A** and based on the following equation:

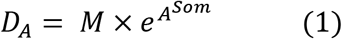

where *D*_*A*_ is the probability of dying of physiological causes <u>at</u> age *A, M* is mutation rate, and *Som* is a composite parameter that determines SMP efficiency. The cumulative distribution function of *D*_*A*_, or the probability of dying of physiological causes <u>by</u> age *A*, resembles human mortality (**Fig. 3B**). The equation should thus provide a robust model for aging-related mortality, reflecting the extended period of high fitness and the late-life accelerating mortality. **Fig. 3A** also demonstrates the relative effects of MR, which is a linear contributor, and the *Som* parameter, which stands for the total damage buffering capacity of the SMP (for details and theory see **Methods: The somatic maintenance program paradigm**). It is important to keep in mind that the *M* parameter (mutation rate) in **Eq. 1** is responsible for the somatic costs of MR (higher MR in **Fig. 3A** accelerates aging-related mortality). Improved SMP, just as body size or other trait, may come at a cost on a short evolutionary time scale, which is later diminished by further adaptation. We did not include this cost in the modeling, since if a trait responds to directional selection this means that the benefit outweighs the cost. Since the amount of change of a trait as a result of positive selection in our model is arbitrary (not exactly copying any particular natural species), we can conclude that this amount of change could already incorporate the net benefit minus cost. In other words, if the benefit of an evolutionary change exceeds its cost, then modeling benefit and cost on an arbitrary scale is mathematically equivalent to modeling only benefit. A summary of life history parameters can be found in Table 1.

**Fig. 3.**
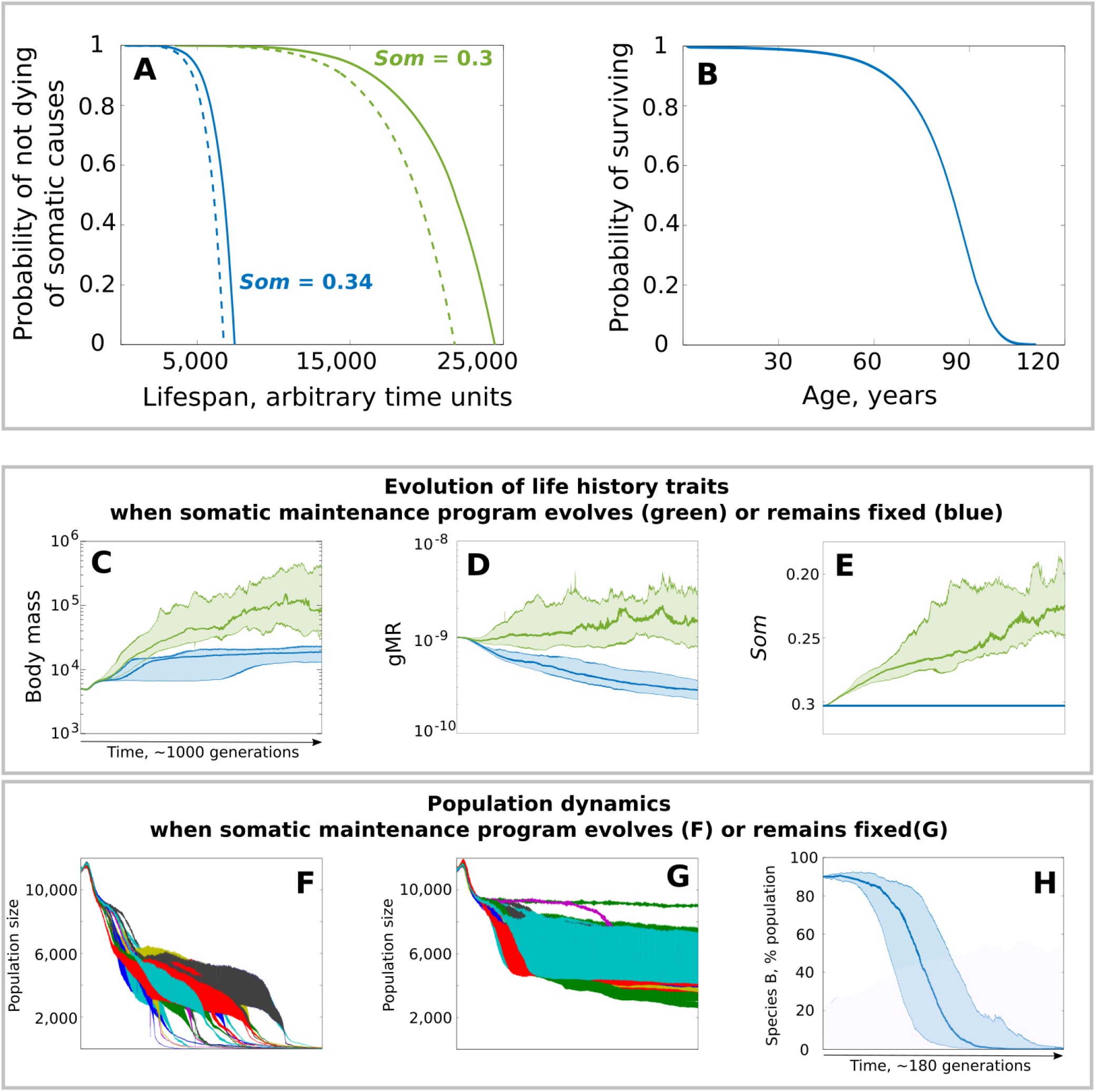
The effect of SMP evolution on the evolution of body mass and mutation rate. (**A**) physiological/aging related mortality curves generated based on the cumulative distribution function of *D_A_* (Eq. 1). Colors represent the effect of the *Som* (SMP) parameter (Eq. 1). Dotted lines were generated by elevating mutation rate 2-fold. (B) modern human mortality in the U.S.A (https://www.ssa.gov). (**C-E**) evolution of life history traits under positive selection for body size. (**F,G**) population size dynamics when SMP can evolve (corresponds to green in *C-E* or SMP evolution is blocked (blue in *C-E);* colors indicate individual populations. (**H**) relative frequency of Species B (SMP evolution blocked, blue in *C-E* in a mixed population with Species A (SMP can evolve, green in *C-E.* For (*C*), (*D*), (*E*) and (*H*) (and similar graphs in other figures), 25 simulations are combined, with the dark line reflecting the mean and shaded area denoting the 95% confidence intervals. The time scale (1,005,000 units) is universal to all results panels (here, C-G) unless otherwise stated.

**Table 1.** A summary of the modeled life history related parameters

| Modeled parameter | Explanation |
| --- | --- |
| Adult body mass | Inherited value of adult body mass. For simplicity, we did not model environmental factors affecting growth and adult body mass |
| Body mass at birth | Inherited body mass at birth |
| Reproductive rate | The frequency of reproduction (time between successive reproductions) |
| Litter size | Inherited litter size (number of progeny per single reproduction) |
| Growth rate | Fixed parameter that was identical for all individuals and invariable |
| SMP | Somatic maintenance program, or an individual's inherited aging profile. Determines the probability of dying of physiological cause as a function of chronological age |
| Age of first reproduction | All individuals started reproducing when their body mass reached 96.93% of their adult inherited body mass. The value was chosen based on initial parameter values so that reproduction started at age 1000 time units. Subsequently the age depended on the evolution of body size and body mass at birth |

### The evolution of SMP promotes selection for increased body size through better tolerance of MR

In our simulations, positive selection for increased body size (**Fig. 3C, green**) led to a concurrent selection for elevated gMR (**Fig. 3D, green**) and improved SMP (**Fig. 3E, green**). Artificially blocking SMP evolution by fixing SMP at the initial value (**Fig. 3E, blue**) significantly slowed the evolution of body size (**Fig. 3C, blue; p ≪ 0.001**) and triggered selection for lower gMR (**Fig. 3D, blue**). We implemented the ecosystem carrying capacity by setting a maximum biomass for the population; therefore, increasing body size led to a corresponding decline in population numbers, amplifying the power of drift (**Fig. 3F,G**). When SMP was allowed to evolve, however, the population entered a “drift zone” when its size decreased to ~4,000 individuals, which shortly thereafter was overcome by selection for even larger body size, visible also by a continuing decline in population numbers (**Fig. 3F**). When we artificially blocked the evolution of SMP, however, the drift zone was more profound. It occurred earlier at the population size of ~6,000-7,000 individuals, and the population was not able to escape from it (for ~1,000 generations) and restore its initial rates of evolution (**Fig. 3G**), indicating an important role of SMP evolution in maintaining evolvability. We further generated a population with two simulated genotypes - Genotype A that could evolve SMP (10% of the population) and Genotype B with SMP fixed at the initial value (90% of the population). We set a maximum population size and removed the maximum biomass limit to rule out body mass effects on population size and selection, and tracked Genotype A and Genotype B frequencies under positive selection for increased body size (for code see **Supplements: Section 1b**). Despite the initial abundance, Genotype B (with fixed SMP) lost the competition in less than 200 generations, reflecting a direct competitive advantage of the capacity to evolve enhanced SMP (**Fig. 3H**). Hereafter, we will call the setting with positive selection for increased body size and freely evolving SMP and gMR the *standard condition* (usually shown in green, unless otherwise indicated) used in comparisons with other selection regimens.

### Abrogating selection for increased body size reduces selection for gMR and SMP

In the absence of positive selection for increased body mass (**Fig. 4A, blue**), both gMR (**Fig. 4B, blue**) and SMP (**Fig. 4C, blue**) demonstrate early positive selection, which appeared to have been caused by rapid evolution of reproductive parameters (see **Supplement: Section 2**). Overall, gMR demonstrates a significant general decrease [non-overlapping confidence intervals (CIs) at the beginning relative to the end of the simulation], and SMP undergoes a significantly smaller improvement compared to the standard condition (green; p ≪ 0.001). Blocking the evolution of body mass (**Fig. 4D, blue**) and SMP (**Fig. 4F, blue**) expectedly led to the evolution of lower gMR (**Fig. 4E, blue**) compared to the standard condition (p ≪ 0.001), which we interpret as being driven by the sMR costs in the absence of benefits of high gMR. In other words, mutation rate is selected against because of its somatic costs and the absence of benefits of higher gMR in static conditions. In natural populations that are under stabilizing selection, gMR will have costs due to greater phenotypic variance from a well-adapted state that are independent of sMR.

**Fig. 4.**
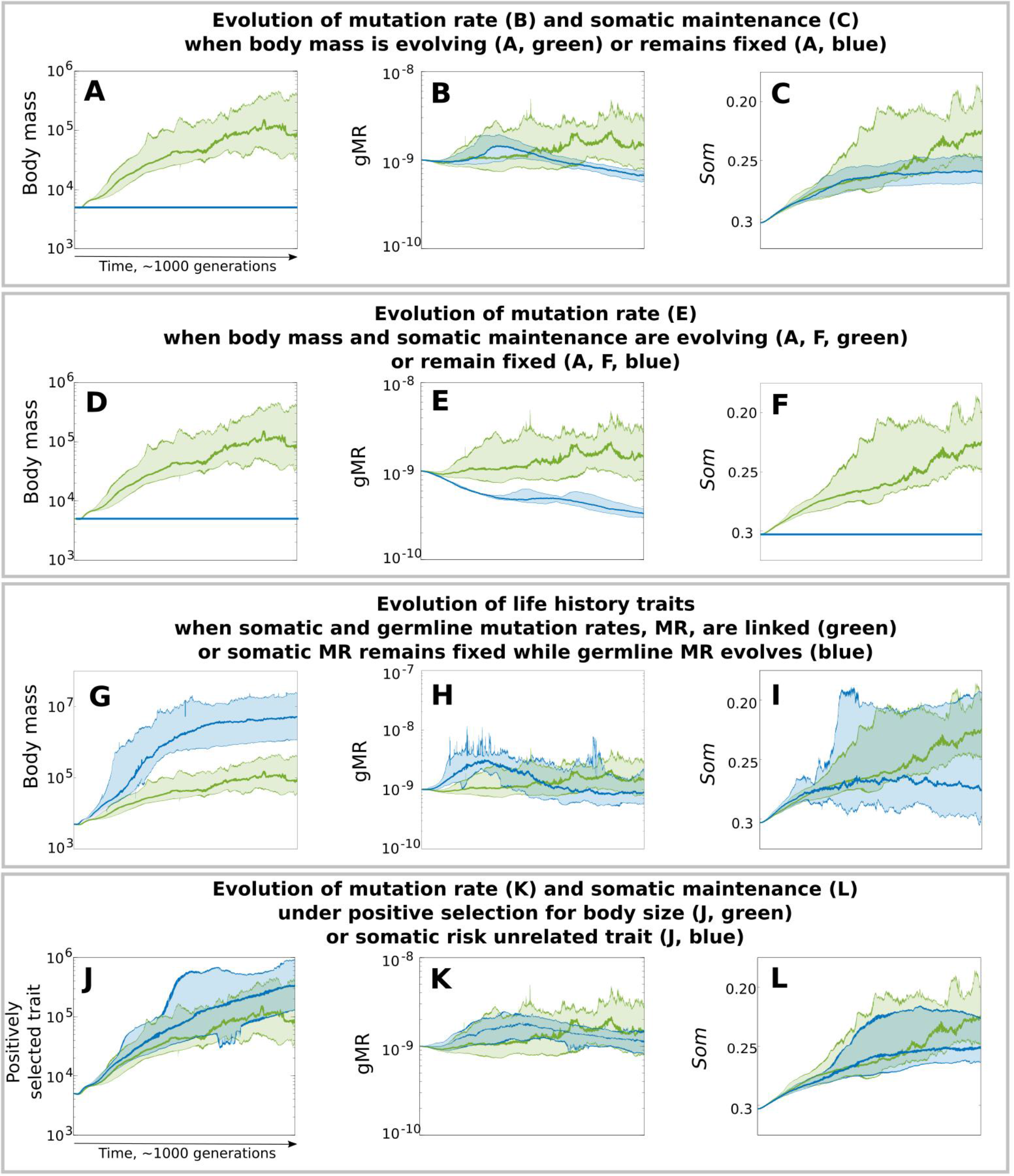
Evolution of body mass, gMR and SMP under various regimens of selection. Separate experiments are stacked as indicated in their subtitles. The layout: left - body size, middle - gMR, right - SMP (the *Som* parameter in Eq. 1) is maintained as in Fig. 3C-E. Green - the standard condition (as green in Fig. 3C-E); blue - alternative conditions with fixed values of a trait (blue horizontal line in *A,D,F*), when gMR and sMR are dislinked so that thesomatic cost is fixed while gMR can evolve (blue in G-I), and under selection for a somatic risk unrelated trait (blue in *J-L*).

### Decoupling sMR cost from gMR enhances the evolution of larger bodies

To investigate the role of the putative gMR benefit versus sMR cost balance in evolution, we further decoupled gMR and sMR by allowing gMR to evolve but making sMR cost fixed and independent of gMR (see **Methods: Model variations**). Decoupling sMR cost from gMR significantly accelerated the evolution of body size (**Fig. 4G, blue**) relative to the standard condition (green; p = 0.0052), revealing that sMR costs can limit the evolution of larger body size. During the early fast evolution of body mass, gMR (**Fig. 4H, blue**) and SMP (**Fig. 4I, blue**) demonstrate a corresponding positive response. Later, further body mass evolution becomes impeded (likely because of the severe depletion in population numbers), coinciding with the evolution of lower gMR. SMP plateaus during this second phase at a significantly lower level compared to the standard condition (p ≪ 0. 001), indicating that the somatic costs of mutation rate stimulate the evolution of more robust SMP.

### Selection acting on a somatic cost-unrelated trait still promotes the evolution of increased gMR and SMP

As we have seen under blocked selection for increased body size (**Fig. 4B,C, blue**), SMP demonstrates an early phase of positive selection (**Fig. 4C, blue**) that is apparently reflected in a corresponding positive selection for higher gMR (**Fig. 4B, blue**). This observation suggests that both SMP and gMR may also respond to selection acting on some other traits, e.g. reproductive parameters (**Supplements: Section 2**). This raises the question whether SMP and gMR evolution would be sensitive to strong selection for a trait that does not affect somatic risks (greater body size increases the target size for somatic mutations). We simulated a condition that was similar to the standard condition, except positive selection was applied to a trait that did not affect sMR related somatic costs (see **Methods: Model variations**); e.g. if SMP improvement is solely a response to the increased sMR cost imposed by larger body, selection acting on an sMR cost-unrelated trait should not drive improvements in SMP. As shown in **Fig. 4J (blue)**, unimpeded by increased sMR costs and declining population size, the evolution of an sMR cost unrelated trait is significantly faster compared to the evolution of increased body size (p ≪ 0.001). Interestingly, gMR (**Fig. 4K, blue**) also demonstrated an early phase of positive selection during early rapid evolution of the selected trait and remains above the initial gMR throughout the entire simulation. As anticipated, SMP is positively selected; however, in the absence of an increasing sMR cost associated with larger bodies, SMP’s improvement is significantly smaller (**Fig. 4L, blue**, p ≪ 0.001). Notably, even with much less enhanced SMP, gMR is still under positive selection in response to positive selection for the sMR cost unrelated trait (**Fig. 4K, blue**), consistent with the sMR/gMR cost/benefit ratio being an important factor regulating selection acting on gMR. Regardless, the results demonstrate that both gMR and SMP are responsive to selection for somatic risk unrelated traits, which indicates that high mutation rate is beneficial in positively selective conditions.

### SMP enables maintenance of gMR when directional selection is weak

As we have seen in **Fig. 4D-F**, in the absence of strong positive selection for increased body size and SMP efficiency, selection acts to lower gMR. **Fig. 5** shows, however, that this selection is significantly modified by the efficiency of SMP. Stronger SMPs (lower *Som* value) relax selection for lower gMR when directional selection is weak (non-overlapping CIs between the standard (red) and either of the improved SMPs). If SMP is strong enough, early evolution for higher mutation rate is observed (Fig. 5, green and blue curves); as we explained this occurs because other traits such as reproductive parameters are evolving during the simulation for some period of time until they reach stasis (as previously shown in Fig. S1). After the stasis was reached, lower gMR is selected for independent of the strength of SMP, as it no longer provides any selective benefit. As will be explained further below, this observation may have significant implication on long-term species survival in relatively static environments.

**Fig. 5.**
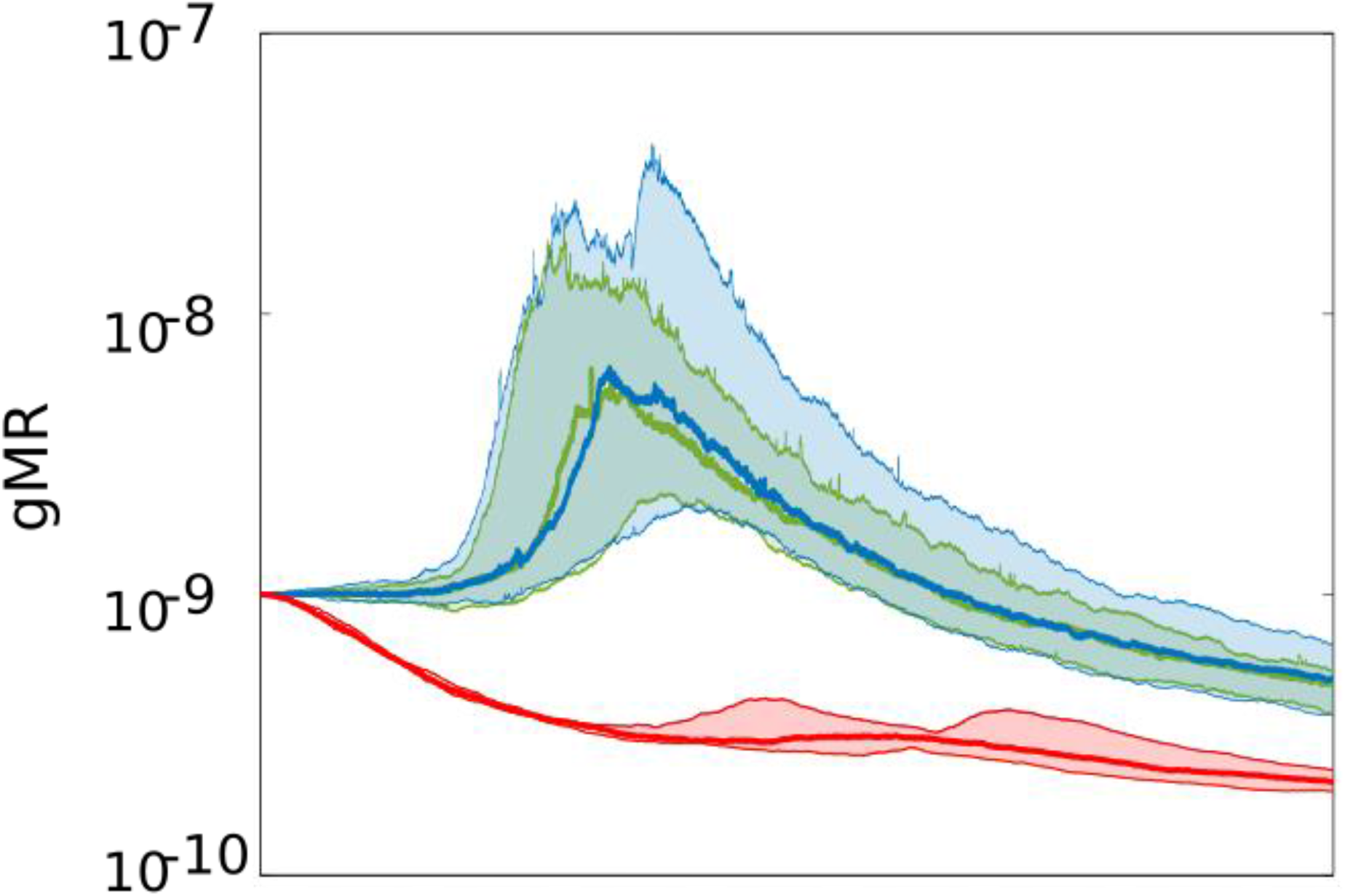
The evolution of gMR in in the absence of positive selection for body mass and SMP. The SMP’s *Som* parameter was fixed at 0.34 (red), 0.24 (green; enhanced 10X) and 0.2 (blue; enhanced 40X); a linear decrease in the *Som* value results in a substantially improved SMP, so that the green SMP is ~10X more efficient compared to red, and the blue is a ~4X more efficient SMP than the green. The standard (red) SMP leads to a significantly stronger selection for lower gMR (non-overlapping 95% CIs); however, the absence of difference between the 10X (green) and 40X (blue) improved SMPs indicates that overly improved SMPs might not provide any further difference for how selection acts on gMR. Note also that even with enhanced SMP (green and blue) there is significant decrease (non-overlapping 95% CIs) in gMR relative to either the initial gMR or to the gMR at the peak (resulting for selection to optimize other traits).

### Modeling competition between a wildtype and mutator phenotypes

Under strong positive selection, whether for increased body mass (**Fig. 3A-C, blue**) or a sMR cost unrelated trait (**Fig. 4H,I, blue**, and **Fig. 4K,L, blue**), we observed consistent signs of positive selection for higher gMR. However, because gMR and sMR are linked, and gMR also confers a cost at the organismal level by increasing phenotypic variance, higher gMR is a trait that should negatively impact individual fitness and therefore be under negative selection. To investigate this question, we mixed two simulated genotypes, one “wild-type” (50%) and one “mutator” (50%) in a population of stable size and under positive selection for a sMR cost unrelated trait. We then observed the genotypes’ frequencies in the population using varying strength of mutators. **Fig. 6A** demonstrates that while the mutator’s fitness initially is lower compared to wild-type, eventually the mutator outcompetes its wild-type counterpart. Interestingly, with increased mutation rate, the magnitude of the mutator’s initial decline increases, but so does the speed at which it subsequently overtakes the population. This result provides a clue for how higher mutation rate, being a trait with negative impact on fitness, can be selected for. Because net organismal fitness is a composite trait impacted by the fitness value of many individual traits, the initial fitness of the “mutator” is lower because, all other traits equal, higher MR incurs increased sMR cost. However, in response to selection, a mutator is capable of more rapidly developing other (adaptive) traits (**Fig. 6B**) and thus its overall fitness soon becomes higher compared to wild-type.

**Fig. 6.**
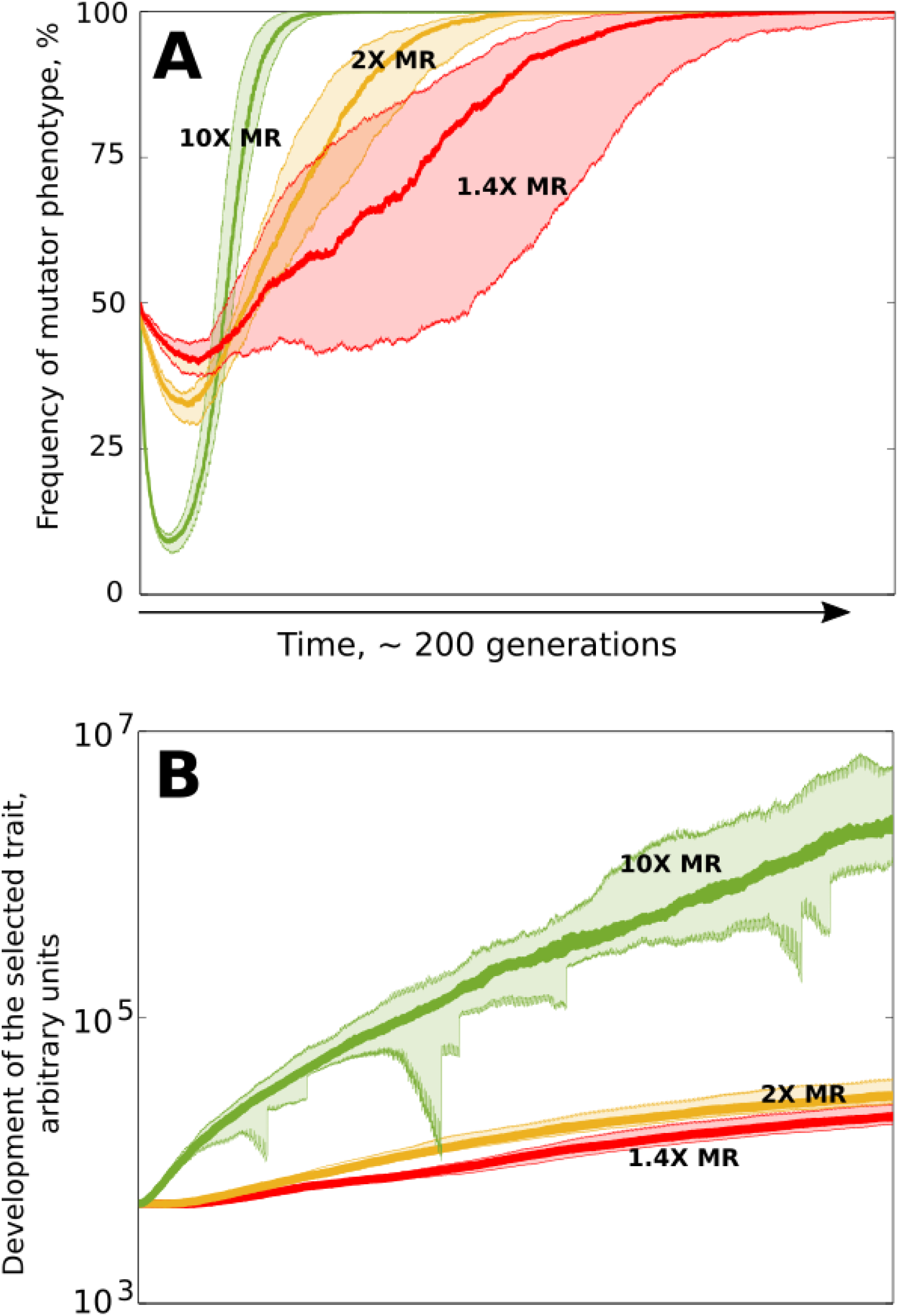
Positive selection for mutators. (**A**) frequency of a mutator phenotype in a mixed competitive population with “wild-type” species. Red (1.4X), orange (2X) and green (10X) are mutators of different fold increase in MR relative to the competitor as indicated by the respective numbers. (**B**) positive selection for a somatic cost neutral trait demonstrates faster evolution (and so adaptation) of mutators. Colors and MR fold increase as in (*A*).

## Discussion

Our study demonstrates that positive selection for increased body size triggers a concurrent selection for improved somatic maintenance to mitigate the increased somatic risks of larger bodies. Improved somatic maintenance, in turn, promotes the evolution of higher germline mutation rates by reducing the cost of somatic mutations and thus altering the sMR/gMR cost/benefit ratio. Conditions of strong positive selection for somatic cost independent traits, as our model shows, can also alter this balance by elevating the benefits of higher gMR. Under stable conditions, alternatively, the sMR/gMR cost/benefit balance is altered by the existing cost of somatic mutations and by the increased cost and absent/reduced benefits of gMR itself, which ultimately favors lower mutations rates. Under stasis, gMR exerts a cost independent of somatic risks by increasing deviation of progeny phenotypes from population mean/median and thus reducing their fitness. Our study thus demonstrates that the evolution of mutation rate is not under a universal population size-dependent selection acting to lower it, but is highly tunable and governed by selection acting on other traits. Importantly, our modeling indicates that under certain conditions elevated mutation rate, unlike perhaps any other trait, can be positively selected despite its negative effects on individual fitness.

Mutation rate in eukaryotes is a highly polygenic trait encoded by multiple genes involved in DNA replication, repair, damage avoidance, and cell division machineries [12,14]. Animals mostly reproduce sexually, which should generate an extensive population allelic diversity for these genes. This diversity should provide for a relatively continuous distribution of mutation rate in populations, rather than being a uniform trait marked with sporadic monogenic mutants, as may occur in asexual populations [24–26]. Such intra-population variation [27,28], as well as the ability of mutation rate to rapidly evolve [29], has been shown for humans. However, sexual reproduction would be supposed to effectively segregate alleles contributing to mutation rate from alleles for other (e.g. adaptive) traits. It has been argued based on other evidence that the efficiency of such segregation in sexual populations is limited [30]. In particular, as we have argued in **Theoretical introduction to the modeling** and shown in Fig.2A-C, the multi-genic nature of the gMR trait should substantially reduce the effects of genetic recombination on the evolution of the net gMR phenotype and its segregation from other traits.

It also appears from our results that animal evolution, with the macroscopic trend toward larger bodies, should have driven a concurrent evolution of extended longevity, the latter being determined by the efficiency of species-specific somatic maintenance programs. Even though extended longevity tentatively appears to be a benefit on its own, e.g. due to extended reproduction period, our model demonstrates that somatic maintenance (and thus longevity) is under a much weaker positive selection in the absence of other positively selected traits. This observation can explain why extended longevity demonstrates significant deviations across animal taxa from the general rule larger body → longer lifespan. Our results indicate that the evolution of longevity (as a function of somatic maintenance efficiency) should be greatly impacted by the rate of evolution of other traits, and not necessarily body size. Of course, longevity will also be highly regulated by external hazards and other extrinsic factors influencing survival probabilities, which has substantial experimental and modeling support [31], which were not explored in our study.

Interestingly, our study predicts an important evolutionary role for the mechanisms of somatic maintenance in addition to their evolution as a means of improving individual survival of large animals [16,21]. Our results demonstrate that selection for enhanced somatic maintenance goes well beyond the evolution of body size and is promoted by strong directional selection acting on any trait. This result indicates that SMPs may have had an important role in the evolution of large animals. Selection for higher gMR following improved SMP may be an important mechanism “rescuing” the reduced evolvability imposed by reduced population size, extended generation times and lower reproduction rates. Therefore, SMPs and longevity may have an important contribution to species’ long-term survival. For example, a prolonged evolutionary stasis [32–35] should trigger selection for lower mutation rates. By relaxing selection for lower mutation rate and thus maintaining evolvability (as shown in **Fig. 5**), enhanced SMPs can ensure better survival of animal groups facing rapid evolutionary transitions or drastically changed environments after such relatively static periods. All other traits equal, species with extended longevity may survive such transitions with higher probabilities.

Lynch and colleagues have provided extensive arguments supporting the idea that the higher MRs in animals compared to unicellular organisms are likely to be caused by reduced population sizes that limit the ability of selection to lower mutation rate [9–11]. In conjunction with population size, in large animals the strength of selection will be further attenuated by lower reproduction rates and extended generation times. Based on our results, Lynch’s theory can be extended by recognizing that somatic maintenance programs (and longevity) should have substantial influence on the general relationship between population size and mutation rates, and on the strength and directionality of selection acting on mutation rates. For example, in our simulation, populations of the same initial size but with different SMP efficiencies demonstrate profound differences in the effects of population size-driven weakening of selection (**Fig. 3F,G**, as well as discrepant selection for mutation rates (**Fig. 3D**).

Selection for higher mutation rates has been shown experimentally in bacteria [24–26,36], whereby engineered or spontaneous mutants with higher mutation rates have been shown to have advantages over wild-type under positively selective conditions. The “mutator hitchhiker hypothesis” explains such selection by the higher probability that adaptive mutations will appear in a mutator cell [26]. Once such a mutation occurs, the mutator genotype spreads to fixation by being genetically linked to the adaptive phenotype.

Modeling studies demonstrate that evolution of evolvability, including varying selection on mutation rates, should be possible in sexually reproducing organisms [30,37,38]. Yet robust experimental corroboration of such a possibility appears to be lacking. Higher mutation rate exerts negative fitness effects on the current generation in any species by imposing somatic risks and contributing to aging. However, in changing conditions whereby directional selection prevails, higher germline mutation rates should provide for better survival of progeny, since directional selection favors phenotypic extremes (tails) in the distribution of phenotypes in a population. Such extreme phenotypes are likely to descend from parents with higher mutation rates (relative to population mean) and thus inherit enriched pools of mutator alleles for multiple MR-related genes. While genetic recombination should be effective in separating any given allele from the adaptive alleles (those that are actually selected for), it should be rather ineffective in segregating the entire multi-genic mutation rate trait from the adaptive phenotype (see Fig.2A-C). Under stabilizing selection whereby population mean (non-deviant phenotypes) is favored, a similar mechanism should trigger co-selection for lower mutation rate by repeatedly sampling the least deviating pool of phenotypes in the population.

In conclusion, our results raise the question of whether the evolution of large body size in animals would be possible without such a complex pattern of selection acting on mutation rate, and whether such a complex relationship is necessary to explain the evolution of large animals. The evolution of large bodies has entailed the cost of losing the ability to evolve via all major parameters that define this ability, such as population size, reproduction rate and generation time, <u>except</u> mutation rate (which increased). Therefore, one scenario could have been that this cost has been so prohibitive for many species that positive selection for mutation rate was necessary to allow evolution of large animals. Alternatively, mutation rate could have been high enough to maintain evolvability at the selection/drift barrier point where selection was no longer able to reduce it further [9]. Understanding which of these scenarios prevails in the evolution of large animals requires more research.

## Methods

### Software

The model was created and all simulations were run in the Matlab environment (MathWorks Inc, MA) version R2014a.

### Model algorithm

The model is a stochastic Monte Carlo type model (the exact algorithm can be found in **Supplements: Section 1a**) that runs a total of 1,005,000 updates (“time” in arbitrary units, AU) unless otherwise stated, which represents ~1000 generations of the simulated animal population (see **Fig. 1** for the flow chart). The simulation starts with building an initial population of 10,000 individuals. Each individual has a number of simulated traits: 1) ID, which is 1 (monogenotypic population) or 1 and 2 (in experiments with competition between two genotypes in a mixed population to indicate genotypes); 2) current age, which increments by 1 at each simulation update; 3) inherited body mass, which is inherited with variation by an individual and will be reached by adulthood (at age ~1000) and equals 5000 AU in the initial population; 4) current body mass, which changes during individual growth, following a growth curve, and plateaus at the inherited body mass in adults; 5) inherited birth mass, which in individuals of the initial population is 300 AU; 6) inherited mutation rate of 10^−9^ AU (explained below); 7) inherited reproduction rate, which is the period with variation between successive reproductions in adult individuals and equals ~600 in the initial population; 8) inherited litter size (initially 1), which is the number of progeny produced per individual per reproduction; 9) inherited parameter of somatic maintenance, which determines the strength of the somatic maintenance program as further explained below; 10) age of first reproduction, which dictates that an individual begins reproducing when its current body mass reaches 0.9693 of its inherited adult body mass (the number is derived so that in the initial population maturity is reached at age ~1000 based on the growth curve).

Each inherited trait varies in progeny relative to parental. This variation was produced by multiplying the inherited mutation rate by the parameter of inherited variance (*inhvar* = 250,000,000) and the product was used as the standard deviation (STD) of the normally distributed variation in inheritance. This transformation was not necessary, as the *inhvar* parameter is constant throughout simulation and it simply determines the magnitude of the mutation rate’s effects in the germline, which is imaginary and in the initial population simply produces 0.000000001 × 25,000,000 = 0.025 that serves as the STD parameter for the normal distribution from which inheritance variation is drawn. However, we kept this two-parametric model for inheritance because mutation rate is also separately used in the equation of the somatic maintenance program (as will be explained later).

Each newborn individual grows, reaches maturity, then reproduces over the rest of its lifetime and eventually dies. The model is asynchronous, so that at every time-point of the simulation the population contains individuals of various ages whose lifecycles develop independently.

And finally, three factors of mortality were modelled in the simulations. First, at every timepoint of the simulation, an individual could die of somatic causes with a certain probability. This probability is small at the beginning of life (but still can be caused by some imaginary inherited genetic defects) and increases exponentially with age based on the paradigm of the aging curve, which is primarily determined by an individual’s inherited somatic maintenance program (SMP). In humans, the aging curve also depends on lifestyle, however we assume in this model that in a wild animal population lifestyle distribution is sufficiently uniform to be neglected. More detailed description of the somatic maintenance paradigm that we applied will be explained further below. Secondly, the simulated animals had a chance of dying of external hazards, such as predators. We applied the Lotka-Volterra model of predator-prey interactions [39,40] to implement the dynamics of predator pressure (effectively the chance of dying of an external hazard cause per timeunit). Here we should mention that smaller individuals and juveniles had higher chances of dying of external hazards, which effectively created positive selection for increased body size and also reflected the typical high mortality rates among juveniles observed in natural populations. And lastly, individuals could die of intra-specific competition. We implemented such competition by setting the upper limit of population’s total biomass, which in nature is imposed by the ecosystem’s carrying capacity. Therefore, in the simulated population biomass produced over the biomass limit caused additional mortality, so that stochastically, population total biomass never exceeded the limit. Larger individuals also had lower probability of dying of intra-specific competition, based on the assumption that competition for resources and mates (the failure to reproduce is effectively an evolutionary death) will typically favor larger individuals and this should have been one of the forces that has been driving the macroscopic animal evolutionary trend towards increasing body size. The advantage of size in this mortality model also created additional positive selective pressure for body size. The total age-dependent mortality of all causes in our model did approximate a typical wild animal mortality curve (**Supplements: Section 3**).

### The somatic maintenance program paradigm

In order to replicate natural mortality caused by physiological aging, such as cancer, decreased immune defense and lower ability to avoid predators or to succeed in intra-specific competition, we made use of the aging curve, or somatic maintenance, concept. Modern humans (in developed nations) and captive animal mortality curves (**Fig. 3B** for human) differ from wild animal mortality curves in very high early life survival with most mortality significantly delayed into advanced ages [41,42]. This difference is caused by many reasons, such as much lower mortality caused by external hazards and better nutrition and general healthcare. It therefore can be assumed that the human and captive animal mortality curves are close representations of the physiological aging curve. As longevity depends on multiple mechanisms of maintaining the soma, we can also call this curve *the somatic maintenance curve*. In order to reconstruct this curve, we assumed that somatic maintenance depends on the interaction of two opposing forces: 1) the accumulation of genetic and structural damage in the soma that promotes aging and 2) the somatic maintenance program consisting of a number of mechanisms that prevent or buffer the effects of genetic and structural damage. The exact mathematical relationship between these two forces and age is not known, however an example of cancer development can be used as a proxy to explain the equation we derived for it.

Oncogenic mutations (including oncogenic epigenetic changes) are the ultimate necessary condition for cancer to develop. The frequency of oncogenic mutations linearly depends on mutation rate on a per cell division basis. Therefore, we assume that linear changes in mutation rate will have linear effects on the odds of the occurrence of oncogenic mutations. An oncogenic mutation provides the initiated cells with a linear change in their fitness relative to normal cells. However, over time an advantageous clone with a constant linear fitness advantage will proliferate exponentially. Therefore, we can already assume that mutation rate should have a linear effect on the cancer curve, while time/age adds an exponential component revealed in an exponential growth of a tumor. We can reasonably assume further that a strong SMP will efficiently suppress such a clone, slowing or even preventing its growth [43]. A weaker SMP will allow the clone to proliferate faster. Therefore, SMP strength can modulate the effects of mutations and time on cancer risk. The exact relationship between SMP strength and physiological risk factors is not known. However, we know that their interaction leads to a net exponent in physiological decline and disease risk.

We therefore reconstructed the human aging curve by maintaining the general principal relationship between these factors as shown in **Eq. 1**. As seen from the equation, mutation rate is a linear contributor to aging. Age itself contributes exponentially, and the somatic maintenance composite parameter *Som* is, in turn, in power relationship to age. The cumulative distribution function of *D*_*A*_ (**Eq. 1**) produces *D(A)* - the probability of dying of somatic/physiological causes <u>by</u> age *A* and yields a shape close to the human mortality curve (**Fig. 3A,B**). We cannot claim that these three factors are in the exact relationship predicted by **Eq. 1**, as it is unknown. As seen in **Fig. 3A**, changes in the *Som* parameter have substantially greater effects on the resulting mortality curve than mutation rate, with mutation rate still having a sizeable effect as well. Yet claims are still made (e.g. [44]) that mutation rate is a larger factor in aging than we assume in this model. Validation of our assumption in general comes from the body of solid evidence that up to 50% of mutations in humans accumulate during body growth by the age 18-20 [45–47]. If mutation accumulation had a significant effect on aging on its own, we should age rapidly until age 18-20 (half-way) and then the rate of aging should decelerate. However, in reality the opposite happens, indicating that the combined strength of the SMP has an overpowering effect in modulating the effects of genetic damage on aging. As a result, we reason that **Eq. 1** might reasonably approximate the natural relationships of these three factors. Therefore, based on an individual’s aging curve we calculated the *D*_*A*_ parameter at each simulation time-point (using the individual’s mutation rate, age and *Som* parameter) and applied it in a binomial trial as the probability of that individual’s dying of somatic/physiological causes in an age-dependent manner. As further explained in **Supplements: Section 4**, the exact relationship between the *Som* parameters and each of the other two (mutation rate and age) has no effect on the model, as the model represents SMP and its variation by using area under the mortality curve. Therefore the sole purpose of **Eq. 1** in the model is to generate an age-dependent curve of physiological mortality whose cumulative function (probability of dying <u>by</u> a certain age) resembles in shape the human mortality/aging curve (see **Supplements: Section 4** for detailed explanation and illustration).

### Model variations

A number of model variations used in simulation experiments are employed. *Fixed trait values* involved simply fixing the initial trait value without inherited variation throughout the entire simulation. *Dislinking of somatic and germline mutation rate* was done by making the value *M* in **Eq. 1** independent of an individual’s mutation rate, which resulted in somatic costs independent of transgenerational variation of mutation rate (effectively from germline mutation rate). *Selection for a trait that did not affect somatic risks* was achieved by transforming the “body mass” trait’s effects by removing the trait from calculations of the risk of death by somatic causes (unlike body size, it did not influence the risk), then removing the population biomass limit and setting maximum population size (unlike body mass, other traits do not directly affect population numbers) and fixing the growth rate curve so that it reached the initial body mass of 5,000 AU (the current body mass parameter in the model; the inherited body mass variation did not exist and the inherited body mass parameter was replaced with the somatic risk unrelated trait). These manipulations made the selected trait a proxy for a trait unrelated to somatic risks (e.g. hair color). *Competitive assays* included individuals with different ID parameters, such as 1 and 2 to indicate different “genotypes”; traits of the “genotypes” then were tracked and stored separately.

### Data processing

Processing of primary data included removal of outliers (see **Supplements: Section 5**). Occasionally the simulations generated “NaN” (not a number) values in individual parameters, which were rare but quickly propagated if left in the population. We immediately deleted individuals from the population if “NaN” values appeared in any of their parameters. Based on the rarity of such events, we can assume that they had the effect of rare early lethal mutations and affected the population at random. Thus, we assume these did not affect the principal results.

### Statistics and data presentation

Most simulation experiments were made with 25 repeats. Due to heavy skews in sample distributions (inferred by D’Agostino-Pearson test for normality of a distribution), all figure panels represent medians (thick lines) and 95 percentiles on each tail (color-shaded areas). Statistical differences between experimental conditions were calculated as follows. We first calculated the sum of all values in each run throughout the entire evolution of a trait (typically 1,005,000 time points). In this way, given the small increment over a long time the sum essentially approximated the area under the curve of a trait’s evolution. These sums (usually 25 repeats in one experiment/sample) were then compared by applying the Matlab implementation of the Wilcoxon rank sum test, which is considered equivalent to the Mann-Whitney U-test. P-values <= 0.05 were considered as indicating significant difference.

## Acknowledgments

We would like to thank Mark Johnston, L. Alex Liggett and David Pollock of the University of Colorado, and Robert Gatenby and Andriy Marusyk of the Moffitt Cancer Center for critical review of the manuscript. These studies were supported by National Cancer Institute grant R01CA180175 to J.D.

## Data availability

All data related to this manuscript will be publically available.

